# High conservation priority of range-edge plant populations not matched by habitat protection or research effort

**DOI:** 10.1101/682823

**Authors:** P. Caissy, S. Klemet-N’Guessan, R. Jackiw, C.G. Eckert, A. L. Hargreaves

**Affiliations:** Department of Biology, McGill University, 1205 Dr Penfield Ave, Montreal QC, Canada, H3A 1B1; Department of Biology, Queen’s University, 116 Barrie St., Kingston ON, Canada, K7L 3N6

**Keywords:** species distributions, range limits, wildlife conservation, cold-edge populations, endangered species, botanical diversity

## Abstract

High-latitude countries tend to contain the polar range-edge of many species that are nationally rare but globally common. This can focus national conservation efforts toward range-edge populations, whose conservation needs and value are disputed. Using plants in Canada as a case study, we ask whether national species-conservation rankings prioritize range-edge populations, and whether conservation priority is matched by habitat protection and research effort. We found that >75% of federally-protected plants only occur in Canada peripherally, at the northernmost 20% or less of their total range, and that the most imperilled taxa had the smallest percentage of their range in Canada (endangered plants: median=1.0%). Occurring peripherally in Canada was associated with higher threat even after accounting for range area, potentially because range-edge taxa experienced 85% higher human population densities in their Canadian range than non-peripheral taxa. High conservation priority was not matched by habitat protection, as more imperilled and more peripheral taxa had smaller fractions of their Canadian range in protected areas. Finally, peer-reviewed research on plants at-risk in Canada was low. Only 42% of plants considered at-risk in Canada had been studied in Canada, and only 11% of species with large distributions outside Canada had been studied in the context of their wider geographic range—information that is critical to establishing their relative conservation value. Our results illustrate that plant conservation in Canada is fundamentally linked to conserving range-edge populations, yet edge populations themselves are understudied, a research gap we must close to improve evidence-based conservation.

## INTRODUCTION

The conservation value of populations at the edge of a species’ distribution is contentious. Species range edges often coincide with declining suitability and abundance of habitat; about three quarters of transplant experiments find declines in performance beyond a species’ range (Hargreaves et al. 2014; Lee-Yaw et al. 2016). Edge populations in poor-quality or isolated habitat are, therefore, predicted to be small (Hengeveld and Haeck 1982; Brown et al. 1996), eroding their genetic quality through genetic drift (Ellstrand and Elam 2003). Small population size and low genetic quality can make edge populations less important to species’ persistence and harder to conserve (Bunnell et al. 2004). However, even dramatically reduced performance beyond the range does not necessarily mean that habitat at the range edge is low quality (Hargreaves and Eckert 2019), and while some edge populations are smaller, less genetically diverse, or more inbred than core populations (Sexton et al. 2011), these patterns are far from universal (Villellas et al. 2013; Pironon et al. 2015, 2017; de Medeiros et al. 2018).

Indeed, range-edge populations may be particularly valuable for species’ long-term success either genetically or geographically. For widespread species, range edges often include extreme or unusual habitats (Thakur et al. 2018); if edge populations are locally adapted, they may contribute uniquely to species’ overall genetic diversity (Bunnell et al. 2004; Sexton et al. 2011). For species whose ranges have expanded and contracted with glacial cycles, populations at the equatorial range edge may harbour disproportionate genetic diversity that could be critical for long-term persistence (Hampe and Petit 2005). Finally, while many taxa are shifting to higher elevations and latitudes in response to climate warming (Chen et al. 2011; Freeman et al. 2018), warming is expected to outpace dispersal ability for 11% of species globally (Thomas et al. 2004). Cool-edge populations are geographically poised to initiate range shifts (Gibson et al. 2009), and may have evolved higher dispersal abilities due to past expansions or demographic instability (Phillips et al. 2010; Hargreaves et al. 2015), making them invaluable for successful range shifts.

The conservation value of range-edge populations is especially germane when conservation policy prioritizes (or deprioritizes) them implicitly or explicitly. Many political jurisdictions rank species by local conservation need, e.g., national, state, or provincial Red Lists. As political borders rarely follow biogeographic boundaries, jurisdictions may contain range-edge populations of species widely distributed outside their borders (hereafter ‘peripheral taxa’; Hunter and Hutchinson 1994). For example, most US states contain >20 peripheral reptiles or amphibians, but many explicitly deprioritize peripheral species in conservation listings, sometimes jeopardizing species’ overall persistence (Steen and Barrett 2015). Conversely, peripheral taxa may be implicitly prioritized; all else being equal they will occupy less area in a jurisdiction, making them more likely to be locally rare and deemed ‘at-risk’ (Lesica and Allendorf 1995; Glass et al. 2017). In jurisdictions with local endemism, local conservation rankings can show considerable mismatch from global conservation priorities. For example, >75% of Finland’s rare beetles (Komonen 2007) and >70% of Canada’s at-risk flora and fauna (Gibson et al. 2009; Cameron and Hargreaves 2020) have wide distributions south of these countries’ borders.

For jurisdictions with many peripheral populations, resolving their conservation value is highly relevant to effective conservation policy, but may require direct study of edge populations themselves. Range position does not consistently predict population size, genetic diversity, or demographic stability (Eckert et al. 2008; Pironon et al. 2017), and the extent of local adaptation to range-edge conditions can be hard to identify without large experiments (Sexton et al. 2011; Hargreaves and Eckert 2019; Anderson & Wadgymar 2020). However, earlier syntheses suggest range edges are under-studied compared to core populations (Eckert et al. 2008; Sexton et al. 2009), potentially impeding evidence-based conservation.

An excellent case study for exploring ‘peripherality’ (the extent to which a species occurs in a jurisdiction at the edge of its range) in conservation is Canada. Canada is the world’s second largest country, spanning almost 10 million km^2^ and >41° of latitude—as much latitude as separates Canada from the Equator—and contains the northernmost potential land for species in the Americas. Canada’s biodiversity is clustered at its southern border (Coristine et al. 2018), as is Canada’s human population, potentially increasing threats to range-edge taxa and creating conflicts between conservation and economic development. Previous estimates suggest ∼75% of terrestrial taxa designated nationally at-risk only occur in Canada at their northern range edge (Yakimowski and Eckert 2007; Gibson et al. 2009), but this has not been formally quantified. Finally, risk-assessments are publicly available for all taxa assessed for federal protection (by the Committee on the Status of Endangered Wildlife in Canada, COSEWIC) and generally include a detailed range map.

We test the relationships between peripherality and: range area in Canada; conservation priority in Canada; conservation risk (human population density) and habitat protection (protected areas); and peer-reviewed conservation research effort. We use vascular plants as they are relatively diverse (3600 species in Canada compared to ∼150 mammals), are of high conservation value globally, and often receive disproportionately little conservation funding (Raven 1987; Schemske et al. 1994). Using COSEWIC risk assessments and range maps (Fig. 1), complemented by NatureServe risk rankings, human population censes, and literature searches, we address five key questions. *Q1)* Do Canadian and global threat rankings differ more for taxa that are peripheral in Canada vs. those that are not, as expected if peripheral taxa have smaller ranges or populations in Canada or face greater threats (see below)? *Q2)* Do more nationally-imperilled taxa have smaller ranges in Canada or smaller proportions of their total range in Canada (i.e. more peripheral), as expected if small ranges and increased peripherality are associated with conservation risk? *Q3*) Do taxa that occur more peripherally in Canada experience higher human population density in their Canadian range, potentially explaining an association between peripherality and conservation risk? *Q4*) Do more-imperilled or more peripheral taxa have a greater proportion of their range in protected areas, potentially indicating active conservation effort? *Q5*) Is conservation research effort evenly spread between range-edge taxa (<20% of their total range in Canada) and non-edge taxa, and how might range-wide studies inform conservation?

**Figure 1.**
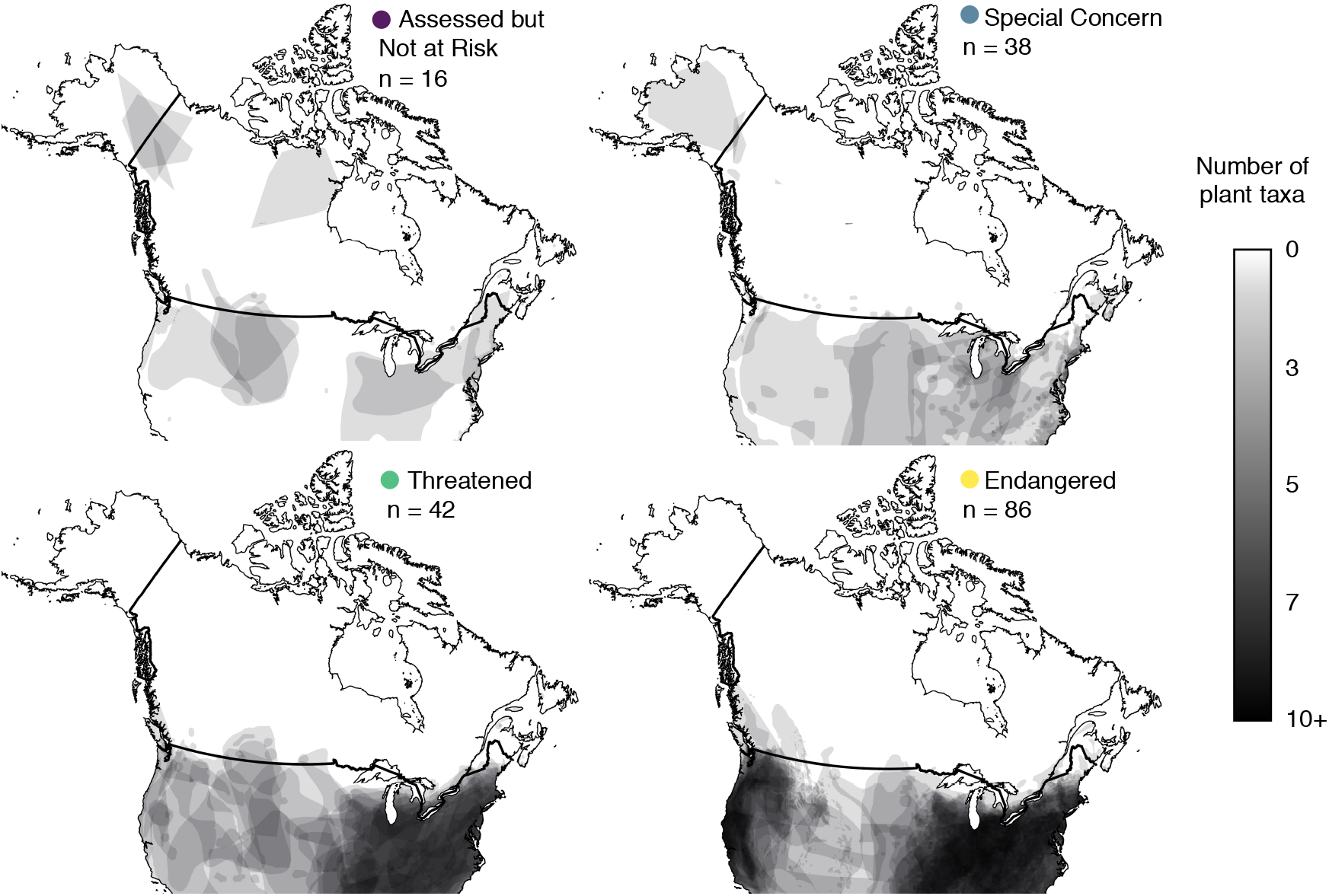
Most plant taxa assessed for protection in Canada only occur in Canada at their northern range edge. Maps illustrate the Canadian distributions, digitized from COSEWIC risk assessments, of 182 of the 220 extant vascular plant taxa assessed from 1977 to 2018, grouped by their COSEWIC-designated risk status (the remaining 38 taxa did not have digitizeable maps but estimates of peripherality are similar; *Table 1*). If taxa were assessed more than once we use the most recent assessment. Most plant taxa in Canada have not been assessed by COSEWIC, so the *Not at Risk* category does not represent all non-imperilled plants but those that were thought to be potentially at risk and later deemed secure. Thick black line shows the Canada-USA border, coloured dots introduce the colour scheme used in subsequent figures.

## METHODS

### Threat status

Flora eligibility for protection under Canada’s Species at Risk Act (SARA) is determined by the COSEWIC Vascular Plant subcommittee using quantitative criteria established by the International Union for the Conservation of Nature (IUCN). COSEWIC was created in 1977, and SARA was passed in 2002 (SARA 2002). There are three steps to a COSEWIC recommendation. 1) COSEWIC prioritizes which taxa to assess. In this phase COSEWIC prioritizes taxa likely to become (globally) extinct (SARA 15.1b). 2) If a taxon is prioritized for assessment, COSEWIC commissions and then reviews and approves an assessment report using the ‘best biological information’ available, including scientific, community, and Aboriginal traditional knowledge (SARA 2002). 3) COSEWIC uses the report to recommend a status: Special concern (may become threatened or endangered due to biological constraints and other threats); Threatened (likely to become endangered if threats not mitigated); Endangered (facing imminent extirpation or extinction), Extirpated, Extinct, or Data deficient (we do not consider Extirpated, Extinct, or Data deficient taxa as they have no current Canadian range or reliable range map); or Not at risk (no imminent risk of extirpation or extinction). COSEWIC determines a status considering only Canadian populations, then reviews whether adjustments are warranted given the likelihood of demographic rescue from populations outside Canada (Environment and Climate Change Canada 2017). The final decision to protect taxa under SARA rests with the federal government after considering the socioeconomic implications of COSEWIC’s recommendation (SARA 2002). We therefore use COSEWIC rather than SARA designations as they more closely reflect biology.

As of August 2018, COSEWIC assessment reports were available for 220 plant populations (1 report each, most available on line but some obtained from COSEWIC directly; sample size details in Table 1). If the taxon had been reassessed we used the most recent assessment. A few taxa had reports for two geographically distinct populations (‘designatable units’ in COSEWIC terminology). If both populations had the same threat status we combined them, otherwise we excluded the taxon as it was unclear how to calculate a corresponding global range per population (Table 1). We excluded reports without range maps or descriptions from which we could estimate the proportion of each range in Canada, yielding 209 taxa including species and subspecies (Table 1). We recorded each taxon’s COSEWIC threat status: Special concern, Threatened, Endangered, or Not at risk; the first three categories are considered ‘at-risk’. COSEWIC only assesses taxa that might be imperilled. The ‘Not at risk’ category does not reflect all plants that are secure in Canada, but a group of taxa deemed potentially at risk (therefore worth assessing) but ultimately secure (e.g., not in sufficient decline for listing, reassessment revealed other populations, taxon not a true taxonomic unit).

**Table 1.** Sample sizes, with data used for each Question in bold. At-risk taxa are those considered Special concern, Threatened, or Endangered, and can have any proportion of their range in Canada. Peripheral taxa include at-risk and not-at-risk taxa. ‘At-risk’ and ‘Peripheral’ columns give the number of populations or taxa and the percentage of the total (i.e. divided by column 2). Analyses for Q2–4 use a continuous measure of peripherality only available for species with digitized range maps. Analyses for Q1 and Q5 define peripherality categorically and so include a larger data set: main analyses use a 20% cut off (species with <20% of their global range in Canada = peripheral), and include taxa with digitized maps and taxa for whom we could estimate peripherality (Y/N) from range descriptions. Using only taxa with digitized maps and a 10% cut-off yielded consistent results (Supp Info).

|  | Extant plants in Canada |  |  |  |  |
| --- | --- | --- | --- | --- | --- |
|  |  |  | Peripheral |  |  |
|  | Total | At-risk<br>(COSEWIC) | 20%<br>cut-off | 10%<br>cut-off | Question |
| (a) populations with a COSEWIC status | 220 | 197<br>(90%) | – | – | – |
| (b) taxa with COSEWIC report and range map<sup>1</sup> | 209 | **189**<br>(90%) | 165<br>(79%)<sup>2</sup> | – | Q5 |
| (c) taxa in (b) with NatureServe ranks for world and Canada | **202** | 186<br>(92%) | 159<br>(79%)<sup>2</sup> | – | Q1 |
| (d) taxa in (b) with digitizeable range map | **182** | 166<br>(91%) | 141<br>(77%)<sup>3</sup> | 127<br>(70%)<sup>3</sup> | Q2, Q3, Q4 |

1. From (a), (b) excludes 2 taxa with no available COSEWIC report (*Hackelia ciliata*, *Symphyotrichum sericeum*), 2 taxa whose COSEWIC report has no map (*Carex nebrascensis*, *Pedicularis furbishia*), and 6 populations from 3 taxa where 2 designatable units (populations) had different COSEWIC statuses (*Eleocharis geniculata*, *Psilocarphus brevissimus*, *Solidago speciosa*). *Smilax rotundifolia* had 2 populations assessed with the same COSEWIC status; these have been merged into 1 taxon. 2. For 27 taxa whose distributions could not be digitized (b minus d), we estimated whether they were peripheral in Canada (20% cut-off) based on undigitizeable maps and range descriptions. We did not do this for the 10% cut-off as it was harder to determine reliably. 3. 27 taxa maps could not be digitized as they did not show the complete range (19), or were in a projection that could not be digitized (4) or were otherwise unclear (4).

For each taxon with a COSEWIC range map, we compiled Canadian and global threat rankings from NatureServe (i.e. ‘rounded global status’; NatureServe 2018). NatureServe ranks use consistent criteria so national and global rankings are directly comparable (a detailed comparison with COSEWIC statuses is in Supp Info Fig. S1). Ranks range from 1 (most threatened) to 5 (least threatened; NatureServe 2018). NatureServe Canadian or global ranks were missing for seven species, so *n* = 202 taxa (Table 1). We calculated the ‘rank disparity’ for each taxon as ‘Global rank – Canadian rank’.

### Range maps & area

Of the 209 COSEWIC range maps, we were able to digitize 182 using the geographic information software *Quantum GIS 2.18* (QGIS Development Team 2018; Fig 1). We georeferenced maps using ≥10 dispersed coordinates of obvious landmarks, such as country, province, county or water-body boundaries. All maps underwent a thin plane spline transformation to allow local deformation, and to standardize map projections to the World Geodetic System 1984 projection (WGS 84, EPSG 4326), a standard projection for worldwide geographic datasets and convenient when working with latitude and longitude coordinates (Tim Elrick, McGill Geographic Information Centre administrator; personal communication). COSEWIC provided maps as polygons (160 taxa) or points occurrences (29 taxa). For polygon maps, we traced digital polygons by hand, and for point occurrence maps, we generated a convex polygon including all points (generally equivalent to COSEWIC’s ‘extent of occupancy’). If we were unable to generate a convex polygon (<4 points on the map; n = 2 taxa), we generated a 1 km buffer around each point creating distinct polygons. We could not digitize maps for 27 taxa whose maps were incomplete or imprecise (as noted by COSEWIC or if mapped as presence/absence by province) or used a projection that did not allow for proper digitization (e.g., circumpolar projection; Table 1).

For each taxon with a range map, we estimated the proportion of its range in Canada. For 182 taxa with a digitized map, we calculated *a)* global range area (km^2^), *b)* Canadian range area (km^2^), *c)* proportion range in Canada (*b* ÷ *a*) as a quantitative measure of peripherality. For area calculations, we projected individual shapefiles in QGIS, using the World Mollweide equal-area projection (WGS 84, EPSG 54 009) since some taxa had large ranges encompassing lower latitudes such as Mexico (Usery and Seong 2000). Range area was extracted with the *gArea* function in the “rgeos” package (version 0.4-1; Bivand and Rundel 2018) in R version 3.2.2. (R Core Team 2017).

For all 209 taxa with COSEWIC range maps, we also assigned a categorical designation of peripherality to simplify analyses comparing national vs. global threat rank and research effort (Questions 1 & 5). We designated taxa as peripheral if they had <20% of their total range in Canada, otherwise not. Any threshold is arbitrary; we chose 20% as all 27 taxa whose map could not be digitized could still be unambiguously categorized as peripheral or not (i.e. clearly had <20% or >20% of their range in Canada) from maps and range descriptions provided in COSEWIC assessments and recovery plans. To test the sensitivity of results we ran two supplementary analyses. First, we re-ran analyses using only taxa with digitized range maps (Table 1d). Second, we used a more conservative cut-off of 10% (Table 1d), the cut-off used by COSEWIC to define their lowest level of Canadian responsibility (Environment and Climate Change Canada 2017).

### Covariates

For taxa with digitized range maps (*n* = 182), we calculated two covariates. First, we estimated the human population density in each taxon’s Canadian range using the dissemination blocks from the 2016 Canadian census (Statistics Canada 2016). Dissemination blocks are the smallest geographic units used by Statistics Canada (equivalent to a city block bounded by intersecting streets, with block size determined by road density), for which inhabitants/block data are available (Statistics Canada 2011). Using *QGIS 2.18* (QGIS Development Team 2018), we overlaid each taxon’s Canadian range polygon on the dissemination block map (McKie 2016). In R, we then summed the inhabitants across all blocks and partial blocks within each species range, and then divided this sum by the taxon’s Canadian range area to estimate human population density.

Second, we calculated the officially protected area within each taxon’s Canadian range using two protected area databases. The Canadian Council on Ecological Areas database includes all levels of protected area (e.g., municipal, provincial, federal) in all Canadian provinces and territories except the province of Quebec (CCEA 2016). Quebec’s ‘Ministère du Développement Durable Environnement et Lutte contre les Changements Climatiques’ database maps all protected areas in Quebec (MDEL 2016). We projected the protected area shapefiles to the World Mollweide equal-area projection (WGS4 EPSG 54 009), then overlaid them on each species’ Canadian range polygon to generate a geographic file of protected habitat. We then calculated the area (km^2^) of protected habitat in each range polygon in R.

### Literature search for peer-reviewed studies on at-risk plants

To assess the peer-reviewed research effort on plants deemed at-risk in Canada, we searched Web of Science for studies on each at-risk taxon with a COSEWIC range map (Table 1b) using its scientific name, English common name, and synonyms listed in its COSEWIC assessment (up to August 2017). We searched all taxa at the species level since few studies were available at lower taxonomic designations (*n* = 209 at-risk species). We narrowed results to ecological or evolutionary studies by including the search term *“*ecolog* OR evolution* OR population* OR demograph* OR genetic* OR conservation* OR fitness”.* We discarded studies that did not present data on the taxon of interest (e.g., only mentioned it in key words), yielding >2900 studies.

We assessed the conservation and geographic relevance of the resulting studies. Studies were deemed conservation relevant if they contained data on natural populations that would be potential use to COSEWIC (e.g., population censuses, performance, life-history, local adaptation, genetic diversity). Studies that contained no data on natural plant populations or data that was not relevant to their conservation (e.g., how much a plant species contributed to a herbivore’s diet) were not considered further. We further classified whether each relevant study sampled wild Canadian populations, wild populations in the USA, both (providing a wider geographic context for at-risk Canadian populations), or neither (sampled populations outside Canada/USA or no specific population). Studies that investigated more than one at-risk taxon (32 studies) were counted for each taxon included.

### Analyses

#### Q1) Do Canadian & global NatureServe ranks differ more for peripheral vs. non-peripheral taxa?

Taxa cannot be less threatened (i.e. more secure) nationally than they are globally, so we are not testing whether a disparity exists or the direction of the disparity, but whether the disparity is bigger for peripheral vs. non-peripheral taxa. Threat-rank disparity (NatureServe Global rank – NatureServe Canadian rank) is numeric but only ranges from 0 (i.e. no difference) to 4 (i.e. taxon ranked as 5 (least threatened) globally and 1 (most threatened) in Canada), requiring non-parametric analyses. We tested whether threat-rank disparity was greater for peripheral vs. non-peripheral taxa using a non-parametric two-sample Wilcoxon test. We used categorical peripherality (peripheral if <20% of range in Canada, otherwise not; Table 1c) to match the (lack of) resolution in the response variable. As detailed above, we tested whether our results were robust to using a) only taxa with digitized maps, and b) defining peripheral as ≥10% of range in Canada (Supp Info).

#### Q2) Do more imperilled taxa have smaller ranges or range proportions in Canada?

We ran one generalized linear model (GLM) for each of three response variables: global range area, Canadian range area, and percentage of range in Canada. Each model considered COSEWIC status as a categorical predictor (model structure: response ∼ status). We predicted that global range area would not differ among COSEWIC ranks, but that Canadian range area and percentage range in Canada would decline with increasing threat. We used negative binomial error distributions for over-dispersed count data (range area), and quasi-binomial distributions for over-dispersed proportional data (range %). Here and for all GLMs, we assessed predictor significance by comparing models with and without the predictor of interest using a likelihood ratio test compared to a Chi-squared distribution (*anova* function in R). When COSEWIC status was significant, we assessed which statuses differed using least squared mean contrasts with a Tukey correction to maintain alpha = 0.05 (package *lsmeans*, version 2.30-0; Lenth 2016). To test whether peripherality was associated with increased imperilment even after accounting for range area, we translated COSEWIC status to integers (1 = Not at risk, 4 = Endangered). We used this ‘numeric status’ as the response in a quasiPoisson GLM, with Canadian range area and proportion of range in Canada as predictors (numeric COSEWIC status ∼ Canadian range area + proportion range in Canada).

COSEWIC procedures changed slightly once SARA was passed in 2002. At the suggestion of people familiar with COSEWIC, we reran all analyses for Q2 including a categorical covariate for whether taxa were assessed before or after 2002, to see whether results differed (Supp Info).

#### Q3) Do more peripheral and/or imperilled taxa have more people in their Canadian range?

We tested whether human population density (response) varied with the proportion of global range in Canada (proportional predictor) and among COSEWIC statuses (categorical predictor), as expected if human population density and peripheral taxa co-occur close to Canada’s southern border (human density ∼ status + proportion range in Canada). We used a negative binomial GLM and assessed significance as for *Q2*.

#### Q4) Do more peripheral and/or imperilled taxa have more of their Canadian range protected?

We tested whether the proportion of a taxon’s Canadian range that overlapped with a protected area (overdispersed proportional response) varied among COSEWIC statuses (categorical predictor) and with peripherality (proportional predictor) using a quasibinomial GLM: proportion Canadian range protected ∼ status + proportion range in Canada. Predictor significance was determined as in *Q2*, and we extracted the fit lines and 95% confidence intervals for the effect of peripherality using the *visreg* R package (version 2.4.1, Breheny and Burchett 2017).

#### Q5) Conservation research effort and insights from range-wide studies on at-risk plants

For the 189 plant species at-risk in Canada with quantifiable ranges (Table 1b), we tested whether peripheral vs. non-peripheral taxa differed in whether or how often they had been studied in the conservation-relevant literature, both in across their entire range and in Canada specifically. All four GLMs used categorical peripherality (Y if <20% of global range in Canada) as a predictor. Response variables were: 1) whether the taxa had been studied anywhere in its range (binomial response/GLM); 2) the number of studies from anywhere in the range (negative binomial response/GLM); 3) whether the taxa had been studied in Canada (binomial GLM); 4) the number of studies that included Canadian populations (negative binomial GLM). The effect of being peripheral was evaluated using likelihood ratio tests, as in *Q2*.

For taxa that have a large fraction of their range outside Canada, studies that sample populations from both Canada and the USA should reveal the most about the relative conservation value of peripheral populations in Canada. We therefore reduced the data above (189 taxa) to species with less than half their range in Canada, selected the conservation-relevant studies on these taxa that included Canadian populations, and read each study carefully to note examples that yield insights relevant to conservation that could not have been gleaned from smaller-scale sampling.

## RESULTS

Of 189 plant taxa considered at-risk in Canada (Table 1b), 152 taxa (80%) occurred in Canada in less than 20% of their range, of which 151 were at their northern range edge (*Micranthes spicata* has the eastern tip of its range in Canada, with the rest in Alaska, Fig. 1 top right). Many at-risk plants occurred only in southern Ontario or British Columbia (68% of 152 peripheral at-risk taxa, 19% of 37 non-peripheral taxa; Fig. 1). Thus, these provinces are disproportionately responsible for conserving at-risk and range-edge plants.

### Q1) Do Canadian & global NatureServe ranks differ more for peripheral vs. non-peripheral taxa?

As predicted, the disparity between NatureServe’s Canadian and global ranks was greater for taxa that are peripheral in Canada compared to taxa that are not (Wilcoxon test, *W* = 5878, *P* < 0.0001, *n* = 202; Fig. 2). Of the 198 plant taxa that NatureServe considered at-risk (ranks 1 to 3) in Canada, 67% were considered secure (ranks 4 or 5) across their global range; most of these nationally-at-risk but globally-secure taxa are peripheral in Canada (123 peripheral, 10 non-peripheral; Fig. 2). Results were consistent using COSEWIC vs. NatureServe Canadian ranks (Fig. S1), or using only taxa with digitized maps and a 10% vs 20% threshold for peripherality (Table S1).

**Figure 2.**
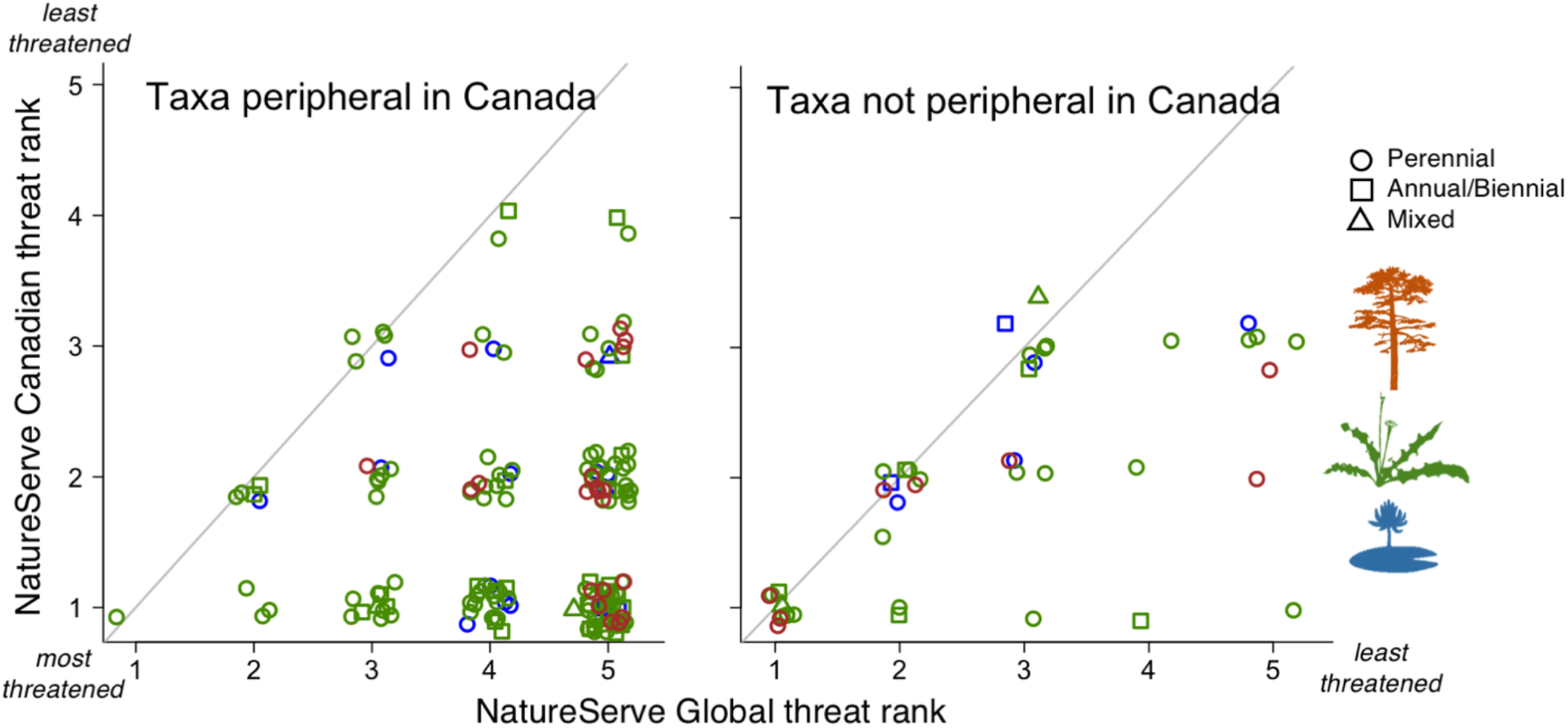
Disparity between Canadian and global threat ranks is greater for peripheral species. (Results for *Question 1*). Diagonal lines indicate Canadian populations have the same threat ranking as the species does globally (values above the line are not possible; points are jittered for visualization). Taxa with <20% of their range in Canada (left, *n* = 159) have a greater mismatch between their Canadian and global threat ranks (more taxa listed as threatened in Canada but secure globally) than taxa with >20% of their range in Canada (right, *n* = 43). Point shape indicates lifespan; colour indicates growth form and habitat (brown = woody shrub or tree, green = non-woody terrestrial plant, blue = aquatic plant). Sample sizes details in Table 1c.

### Q2) Do more imperilled taxa have smaller ranges or range percentages in Canada?

Plants considered at-risk by COSEWIC (status = Special Concern, Threatened, or Endangered) generally had large global distributions (median = 390,521 km^2^), with much smaller ranges in Canada (median = 4,598 km^2^) that were clustered toward Canada’s southern border (Fig. 1). At-risk taxa had a median of 1.8% of their global range in Canada. As predicted, global range size did not differ among Canadian status categories (χ^2^_df3_ = 7.5, *P* = 0.057; Fig. 3a). However, taxa assessed and deemed at-risk had significantly smaller Canadian ranges than taxa assessed and deemed Not-at-risk taxa (χ^2^_df3_ = 40.7, *P* < 0.0001; Fig. 3b), and the most imperilled taxa had the smallest percentages of their range in Canada (χ^2^_df3_ = 15.0, *P* = 0.0002; Fig. 3c). Results did not differ between taxa assessed before or after SARA was passed (Table S2).

**Figure 3.**
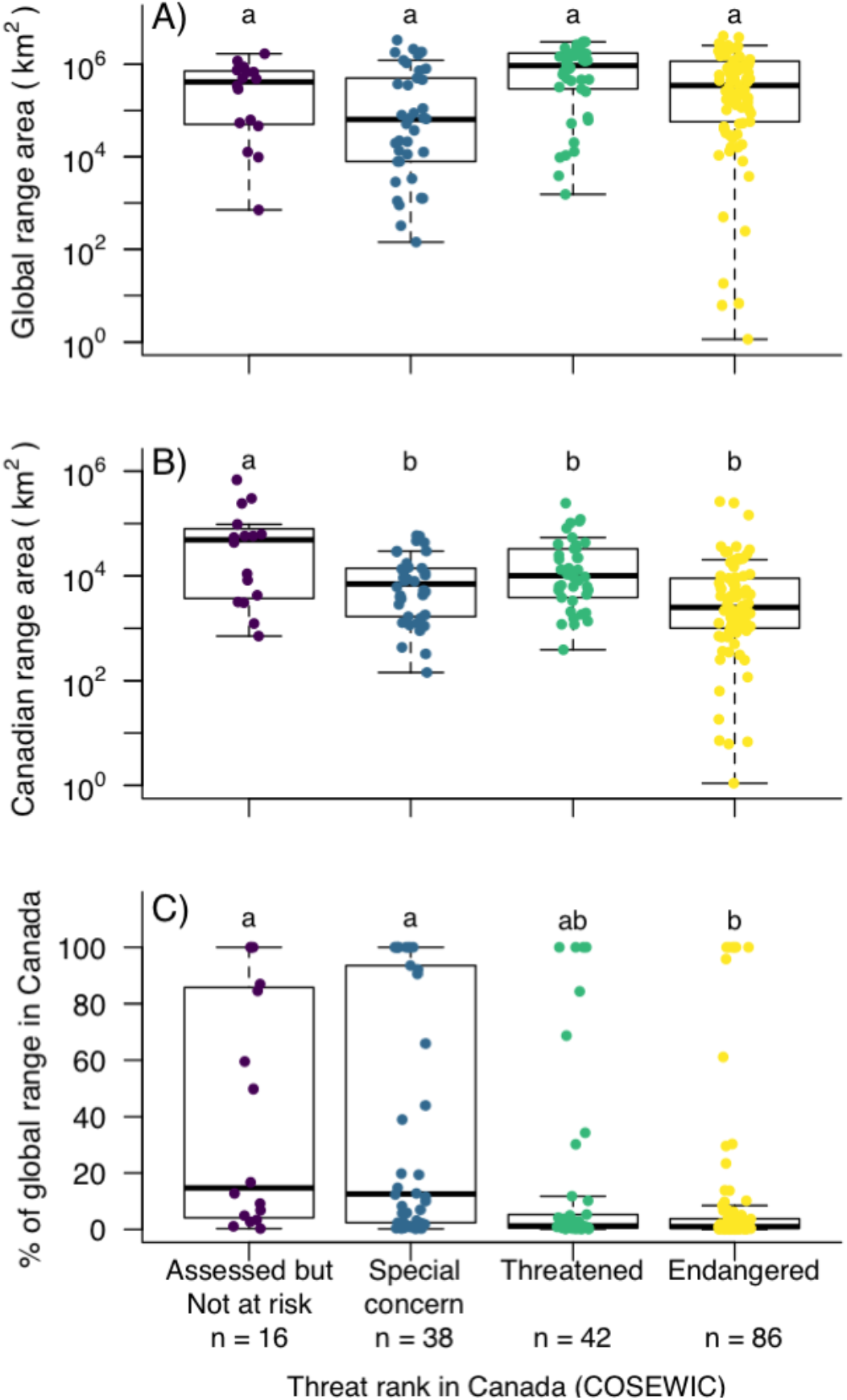
Global vs. Canadian range size for plants assessed for protection in Canada. (Results for *Question 2*). Differing letters indicate significant differences among COSEWIC threat statuses within panels. The *Not at Risk* status does not represent all non-threatened plants but those that were thought to be potentially at risk but later deemed secure. The lower, middle and upper horizontal lines in each boxplot indicate the 25^th^ percentile, the median and the 75th percentile, respectively. Whiskers extend 1.58 interquartile range / square root *n* from the median or to the extreme points, whichever is less. Coloured points show raw data (horizontal jitter to facilitate visualization of overlapping points); sample sizes in Table 1d.

The extent to which taxa occurred in Canada at the edge of their range was associated with increased conservation threat, even after accounting for absolute range area in Canada. Numeric COSEWIC status increased (i.e. more imperilled) as range size in Canada decreased (χ^2^_df1_ = 8.8, *P* = 0.003) and as range percentage in Canada decreased (χ^2^_df1_ = 16.9, *P* < 0.001; Fig. S3).

### Q3) Do more peripheral and/or imperilled taxa have more people in their Canadian range?

Human population density differed with threat status, but not with peripherality. More imperilled plants had significantly higher human population densities in their Canadian ranges compared to taxa assessed at lower threat ranks (χ^2^_df3_ = 36.7, *P* < 0.0001; Fig. 4). After accounting for differences among threat ranks, taxa that are more peripheral in Canada did not have higher human densities within their Canadian range (χ^2^_df1_ = 1.7, *P* = 0.20).

**Figure 4.**
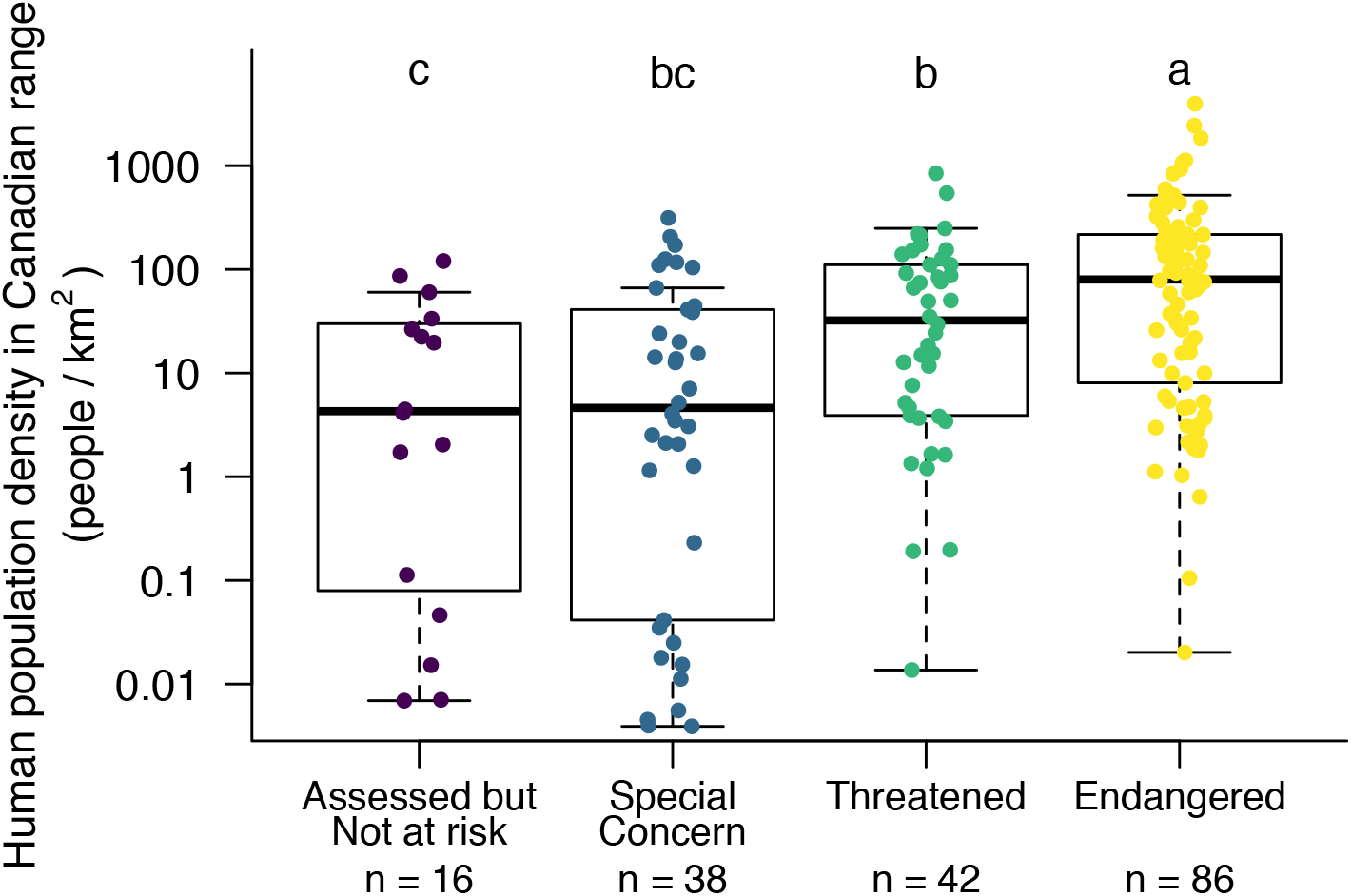
Human population density varied among COSEWIC threat statuses. (Results for *Question 3*). Boxplots are as in Fig. 3. Sample size details in Table 1d.

### Q4) Do more peripheral and/or imperilled taxa have more of their Canadian range protected?

Plant taxa considered at-risk in Canada had only 3.7% (median) of their Canadian range protected, and only 8 taxa had >50% of their Canadian range protected (Fig. 5). Habitat protection varied among threat statuses (COSEWIC status: χ^2^_df3_ = 15.1, *P* = 0.0017), but contrary to predictions the taxa the are the most imperilled in Canada tended to have the lowest fraction of their Canadian range protected; Fig. 5a). Habitat protection also varied with peripherality (proportion range in Canada: χ^2^_df1_ = 37.0, *P* < 0.001). As predicted, the more peripherally a taxon occurred in Canada, the smaller the fraction of its Canadian range was protected (Fig. 5b).

**Figure 5.**
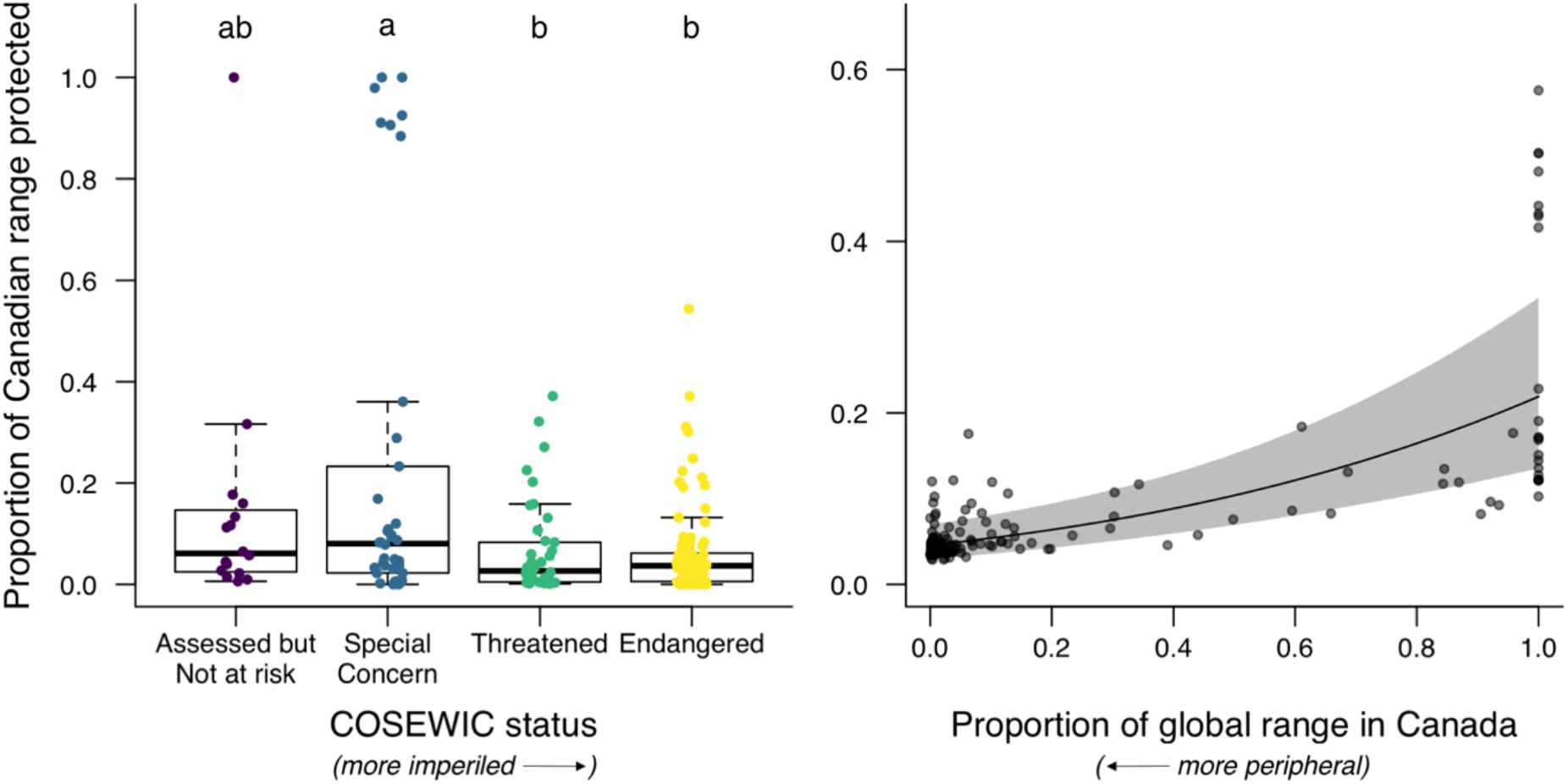
Habitat protection varies with threat status and peripherality. (Results for *Question 4*). (A) Taxa more imperilled in Canada (threatened, endangered) have the smallest proportion of their Canadian range protected. Boxplot formatting and *n* as in Figs. 2&3. (B) Taxa with less of their global range in Canada (more peripheral) have smaller proportions of their Canadian range protected. Line, shading, and points show fit, 95% confidence intervals, and residuals extracted from the quasi binomial GLM: proportion Canadian range protected ∼ COSEWIC status + proportion range in Canada.

### Q5) Conservation research effort and insights from range-wide studies on at-risk plants

Our literature searches yielded 657 peer-reviewed, conservation-relevant studies on the 189 plant species that are at-risk in Canada and whose geographic distribution we can assess (Table 1b). Almost half (44%) of the 189 species had not been studied in peer-reviewed work that could inform conservation (Fig. 6a). Though this does not preclude the existence of studies in the non-refereed literature or journals not indexed on Web of Science, it suggests that the ‘best biological information’ is sparse for many taxa. Compared to species with more than 20% of their range in Canada, species that only occur peripherally in Canada did not differ in the likelihood that they had been studied (χ^2^_df=1_ = 1.73, *P* = 0.19) or in the number of studies (χ^2^_df=1_ = 0.21, *P* = 0.88; Fig. 6a). Comparisons remained non-significant if we used only taxa with digitized range maps or 10% peripheral criterion (Table S3).

**Figure 6.**
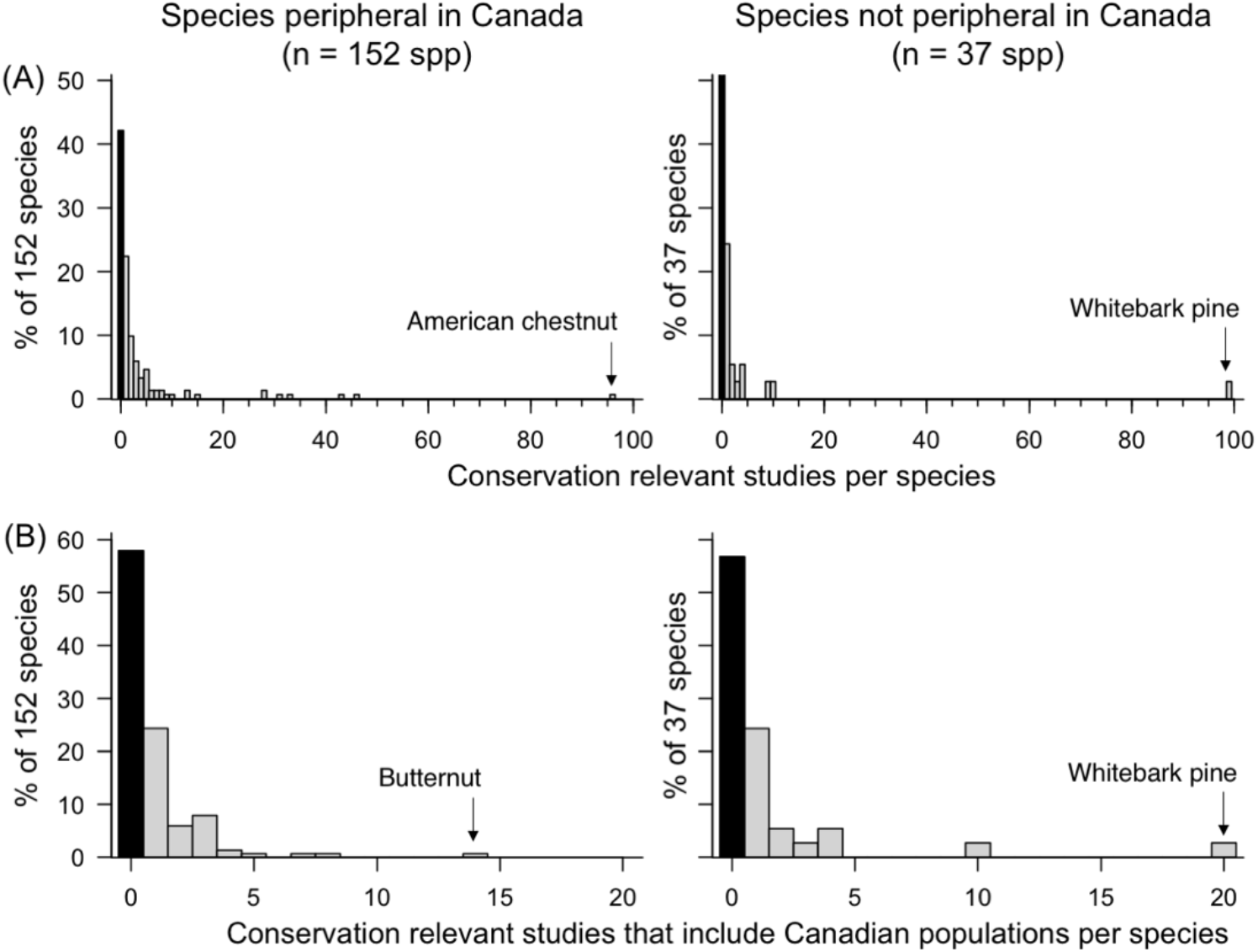
Distribution of conservation-relevant research on plants at-risk in Canada. (A) distribution of all 657 studies that could inform conservation; (B) 187 studies from (A) that included data from wild Canadian populations. Left panels = peripheral species (< 20% of range in Canada); right panels = non-peripheral species. Black bars show species with no peer-reviewed studies.

Of 657 conservation-relevant studies, only 187 included Canadian populations. Less than half (42%) of plant species at-risk in Canada had been studied in Canada (Fig. 6b). Peripheral and non-peripheral species did not differ in the likelihood that they had been studied in Canada (χ^2^_df=1_ = 0.02, *P* = 0.90), nor in the number of studies that included Canadian populations (χ^2^_df=1_ = 2.72, *P* = 0.099; Fig. 6b). However, if one considers only taxa with a digitized range map or a stricter definition of peripheral (<10% range in Canada), peripheral species had fewer studies that included Canadian populations than non-peripheral species (Table S3).

Most (162 of 189) plant species at risk have less than half their range in Canada, such that understanding Canadian populations in the context of their wider geographic range could inform conservation. But only 6% (34 of 536) of studies on these species included both Canadian and USA populations, and these 34 studies covered only 20 (11%) of the 162 species. Studies that performed range-wide sampling provided unique insights into conserving peripheral populations (Table 2). These include whether edge populations differ from core populations genetically, demographically, or in key traits or habitat affinity. For instance, populations of Deerberry (*Vaccinium stamineum*) decreased in size and frequency toward the species northern range edge in Canada, but were nevertheless as productive and genetically diverse as core populations, and showed evidence of local adaptation and high dispersal ability (Yakimowski and Eckert 2007, 2008). Thus the demographic and genetic value of these populations was not predicted by their peripherality, size, or spatial isolation.

**Table 2.** Studies with wide geographic coverage can shed light on the conservation needs and value of peripheral populations. Key findings are taken from studies of plant taxa deemed at-risk in Canada that sampled both Canadian and USA populations. All taxa are peripheral in Canada.

| **Response** |  |  |
| --- | --- | --- |
| Finding for edge populations | Species | Reference |
| **Genetic uniqueness** |  |  |
| Genetically differentiated from core populations (neutral variation) | Branched bartonia | Ciotir et al. 2013 |
|  | Green dragon | Boles et al. 2000 |
|  | Spalding's Campion | Lesica et al. 2016 |
| Greatest population differentiation | Butternut | Hoban et al. 2010 |
| **Genetic diversity** |  |  |
| Not lower neutral genetic diversity than core populations | Heartleaf plantain | Mymudes and Les 1993 |
|  | Deerberry | Yakimowski and Eckert 2008 |
|  | Green dragon | Boles et al. 2000 |
|  | Golden paintbrush | Godt et al. 2005 |
|  | Cucumber tree | Budd et al. 2015 |
| **Population demography** |  |  |
| Not smaller than core populations | Golden paintbrush | Godt et al. 2005 |
| Smaller & more isolated than core populations | Deerberry | Yakimowski and Eckert 2007 |
| **Performance & ecology** |  |  |
| Similar habitat to core populations | Small whorled pogonia orchid | Mehrhoff 1989 |
| Lowest seed viability & distinct dormancy patterns | Green dragon | Yang et al. 1999 |
| Higher seed mass & equal sexual reproduction / productivity as core populations | Deerberry | Yakimowski and Eckert 2007 |

Branched bartonia = *Bartonia paniculata ssp. paniculata*; Butternut = *Juglans cinerea*; Cucumber tree = *Magnolia acuminata*; Deerberry = *Vaccinium stamineum*; Golden paintbrush = *Castilleja levisecta*; Green dragon = *Arisaema dracontium*; North American ginseng = *Panax quinquefolius*; Small whorled pogonia orchid = *Isotria medeoloides*; Heartleaf plantain = *Plantago cordata*; Spalding's campion = *Silene spaldingii*.

## DISCUSSION

Our results show that conservation of plants in Canada is fundamentally the conservation of range-edge populations. Three quarters of nationally at-risk plant taxa only occur in Canada at the northernmost 20% or less of their global range, in line with earlier estimates for all at-risk taxa combined (Gibson et al. 2009). We do not have range maps for the thousands of plants in Canada that have not been assessed by COSEWIC, so cannot directly test whether at-risk plants are more peripheral than average. However, taxa with <20% of their range in Canada had a greater disparity between their Canadian vs. global threat ranking than taxa with more of their range in Canada (Fig. 2), and the most imperilled taxa were significantly more peripheral (smaller proportion of their range in Canada) than less imperilled taxa (Fig. 2C), suggesting a real relationship between occurring as range-edge populations and being nationally at-risk.

Range-edge taxa could be more nationally threatened because they have smaller ranges and therefore fewer individuals in Canada, or because their Canadian populations are disproportionately threatened. Our results suggest that both are true. Smaller range area in Canada was associated with higher COSEWIC threat status (Fig. 3B), but peripherality was associated with higher threat even after accounting for Canadian range area (Fig. S3). Endangered taxa are both the most imperilled and most peripheral group (Fig. 3C) and had significantly higher human population densities in their Canadian range (Fig. 4), although peripherality was not associated with human density after accounting for threat rank. We did not test effects of human activity not associated with high population density (e.g., agriculture), but overall human activity is also highest in southern Canada (Coristine and Kerr 2011), where most-at-risk and almost all peripheral taxa occur (Fig. 1). Thus, higher national threat ranks for peripheral taxa probably reflect real increased risk per population.

Stewarding Canada’s future biodiversity clearly requires an informed policy on conserving peripheral populations. Not only are most at-risk flora range-edge populations, but these populations are geographically poised to initiate northward range shifts under climate warming (Gibson et al. 2009), and so will make up more of Canada’s flora in the future. Unfortunately, conservation of peripheral taxa has been debated in the absence of much relevant scientific evidence. Less than half the plant species with at-risk populations in Canada have been studied in Canada in a way that could guide their conservation. While this could reflect difficulty in obtaining permits or adequate sample sizes, taxonomic bias is likely. For example, one bird species that is both peripheral and at-risk in Canada had almost 50 studies that included Canadian populations (Marbeled murrelet; Web of Science search May 2019), far more than any peripheral at-risk plant species (Fig. 6B).

The few conservation-relevant studies that include both Canadian and US populations illustrate the value of studying peripheral populations directly and in a broad geographic context. However, most of these studies have assessed neutral genetic diversity and population structure (Table 2). Conservation would particularly benefit from studies of characteristics important for long-term persistence and range expansion, such as habitat preferences, population demography and dispersal ability (Schemske et al. 1994). Future genetic work could move beyond neutral variation to evaluating the adaptive diversity likely to be important in responding to environmental change (Shaw and Etterson 2012), and local adaptation through which range-edge populations may contribute uniquely to species’ biodiversity (Yeaman et al. 2016). Whether researchers will close these knowledge gaps depends partially on how government agencies incentivise (i.e. fund) and remove barriers to (i.e. permit) research on at-risk peripheral populations. Unfortunately, the “peripherality issue” is not currently highlighted in federal programs that fund species-at-risk research in Canada (e.g. Government of Canada 2019).

We hope that exposing this research need inspires future work on at-risk edge populations, but recognize that amassing this work will take time, and that some of the most informative types of study, e.g., large reciprocal transplants, will be impossible with endangered taxa. In the meantime, we have a potentially under-used body of research that could inform Canadian conservation: the already extensive theory and empirical research on species range edges (Sexton et al. 2009; Pironon et al. 2017). While this research clearly shows that edge populations can vary significantly from one another in demography (Sagarin et al. 2006) and degree of adaptation (Hargreaves and Eckert 2019), it also reveals broad scale patterns that can be predictive, e.g., that poleward range edges are often dispersal limited whereas high-elevation edge populations are often demographic sinks (Halbritter et al. 2013; Hargreaves et al. 2014), and suggests novel conservation strategies, e.g., increasing gene-flow among isolated range-edge populations to spread broadly beneficial alleles (Sexton et al. 2011; Hargreaves and Eckert 2019). For countries like Canada whose biodiversity is disproportionately comprised of range-edge populations, leveraging this literature could help meet commitments to protecting current and future biodiversity.

## ACKNOWLEDGEMENTS

We thank Christie Whelan (Dept Fisheries and Oceans), Gary Allen & Mark Major (Parks Canada), Joanna Freeland (Trent University), Jeanette Whitton (UBC), and Karen Samis (Queen’s U) for providing insight into Canadian conservation policy and practise. We also thank Tim Elrick (McGill University) for his help with geographical data. Funding was provided by Natural Sciences and Engineering Research Council of Canada (Discovery grants to ALH and CGE), by E. Gordon Edwards and Vivian Astroff Science Undergraduate Research Awards to PC, and by William E. Bembridge Science Undergraduate Research Award to SKN.

## SUPPLEMENTARY INFORMATION

### QUESTION 1

#### Comparability of NatureServe and COSEWIC ranks

Although Nature Conservancy and COSEWIC ranks use different numbers of categories (five and four, respectively), both are derived from IUCN criteria. To test whether the two bodies ranked taxa consistently, we converted COSEWIC ranks to numeric ranks from 4 (not-at-risk) to 1 (endangered); as NatureServe did not give any species a 5 (least threatened) in Canada, both NatureServe Canadian ranks and numeric COSEWIC ranks varied from 1 to 4. We used a paired Wilcoxon test to assess whether COSEWIC and NatureServe ranks for Canadian populations differed overall. NatureServe and COSEWIC ranks did differ significantly, as NatureServe tended to consider taxa more nationally threatened than COSEWIC (Fig. S1).

**Figure S1.**
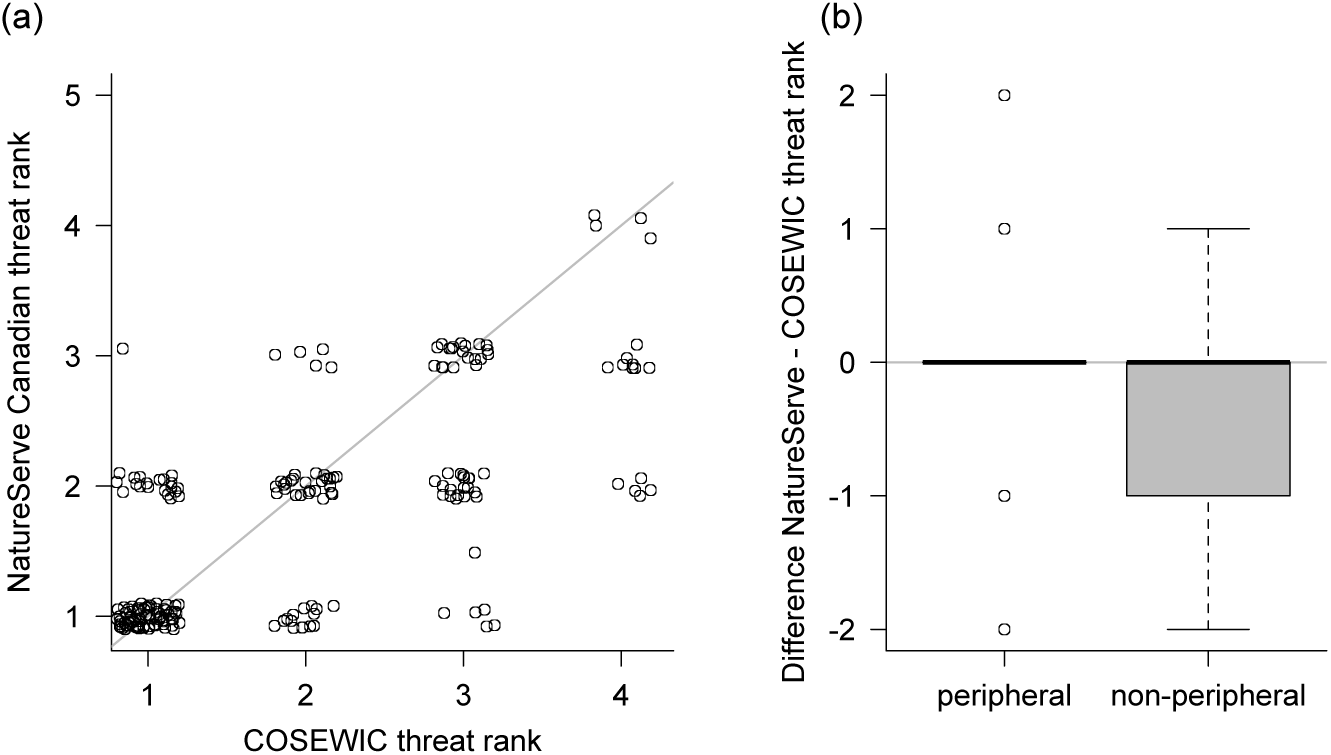
(a) NatureServe tended to consider taxa more threatened than COSEWIC, i.e. more points fall below the diagonal line reference line than above (1 = most threatened; Wilcoxon paired rank test = 924.5, *P* = 0.0003). (b) This difference reflected higher threat rankings for non-peripheral taxa (Wilcoxon unpaired rank test = 4139, *P* = 0.016). Centre lines and boxes show the median, 25th and 75th quartiles (R Core Team 2017).

However, the systematic difference between NatureServe Canadian ranks and COSEWIC ranks did not alter conclusions for Q1. If we used COSEWIC ranks instead of NatureServe Canadian ranks to calculate the discrepancy between global and national conservation priority, peripheral taxa still have greater discrepancy than non-peripheral taxa (Fig. S2).

**Figure S2.**
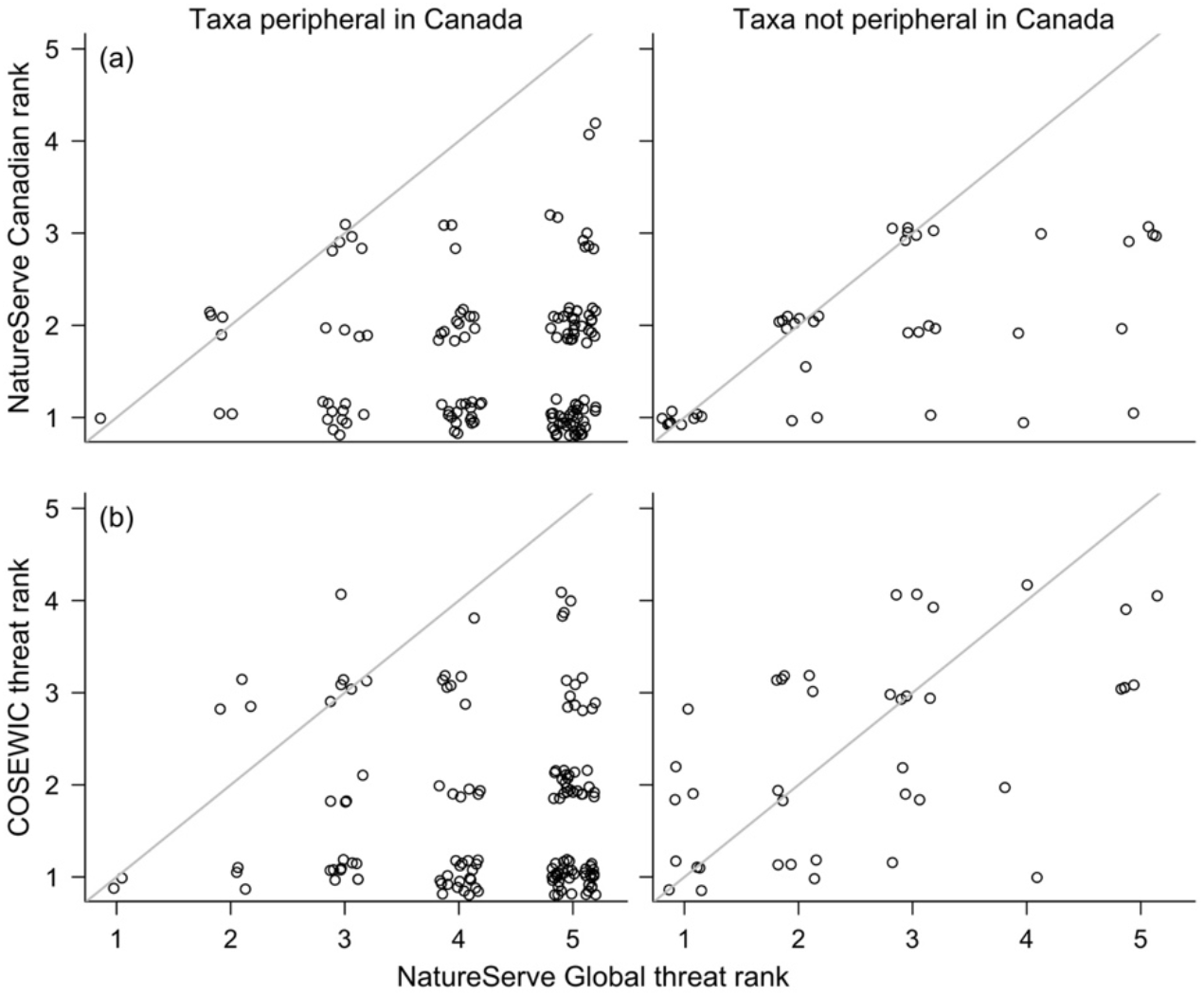
The disparity between national and global threat ranks is greater for species peripheral in Canada, whether one assesses Canadian threat using NatureServe Canadian ranks (shown in (a) – same data as depicted in Fig 2) or COSEWIC ranks (b: Wilcoxon unpaired test on whether the disparity between global and Canadian threat ranks differs between peripheral and non-peripheral taxa: 5877.5, *P* < 0.0001). Diagonal lines indicate Canadian populations have the same threat ranking as the taxon globally. Taxa that are peripheral in Canada (left) have a greater mismatch between their Canadian and global threat ranks (more taxa listed as threatened in Canada but secure globally) than taxa with >20% of their range in Canada (right).

#### Changing definition of peripheral

Our main analyses used 202 taxa (Table 1c) and paired Wilcoxon tests to test disparity between NatureServe Canadian and global rankings among taxa deemed peripheral in Canada (<20% of global range in Canada, determined from digitized maps or maps and range descriptions) or non-peripheral. We tested the sensitivity to our definition of ‘peripheral’ by rerunning analyses using only taxa with a digitized range map and quantitative assessment of peripherality, and using a stricter threshold for ‘peripheral’ of 10% of range in Canada. Conclusions did not change (Table S1).

**Table S1.** Reanalysis of Question 1 with different definitions of peripheral. Top row gives results reported in the main paper.

| Definition of peripheral | *n* taxa |  |
| --- | --- | --- |
|  | total (peripheral) | Wilcoxon *W*, *P* |
| <20% of range in Canada, determined from digitized maps or non-digitized maps & range descriptions | 202 (159) | 5878, *P* < 0.0001 |
| <20% of range in Canada, determined from digitized maps only | 175 (135) | 4686, *P* < 0.0001 |
| ≤10% of range in Canada, determined from digitized maps only | 175 (123) | 684, *P* < 0.0001 |

### QUESTION 2

**Figure S3.**
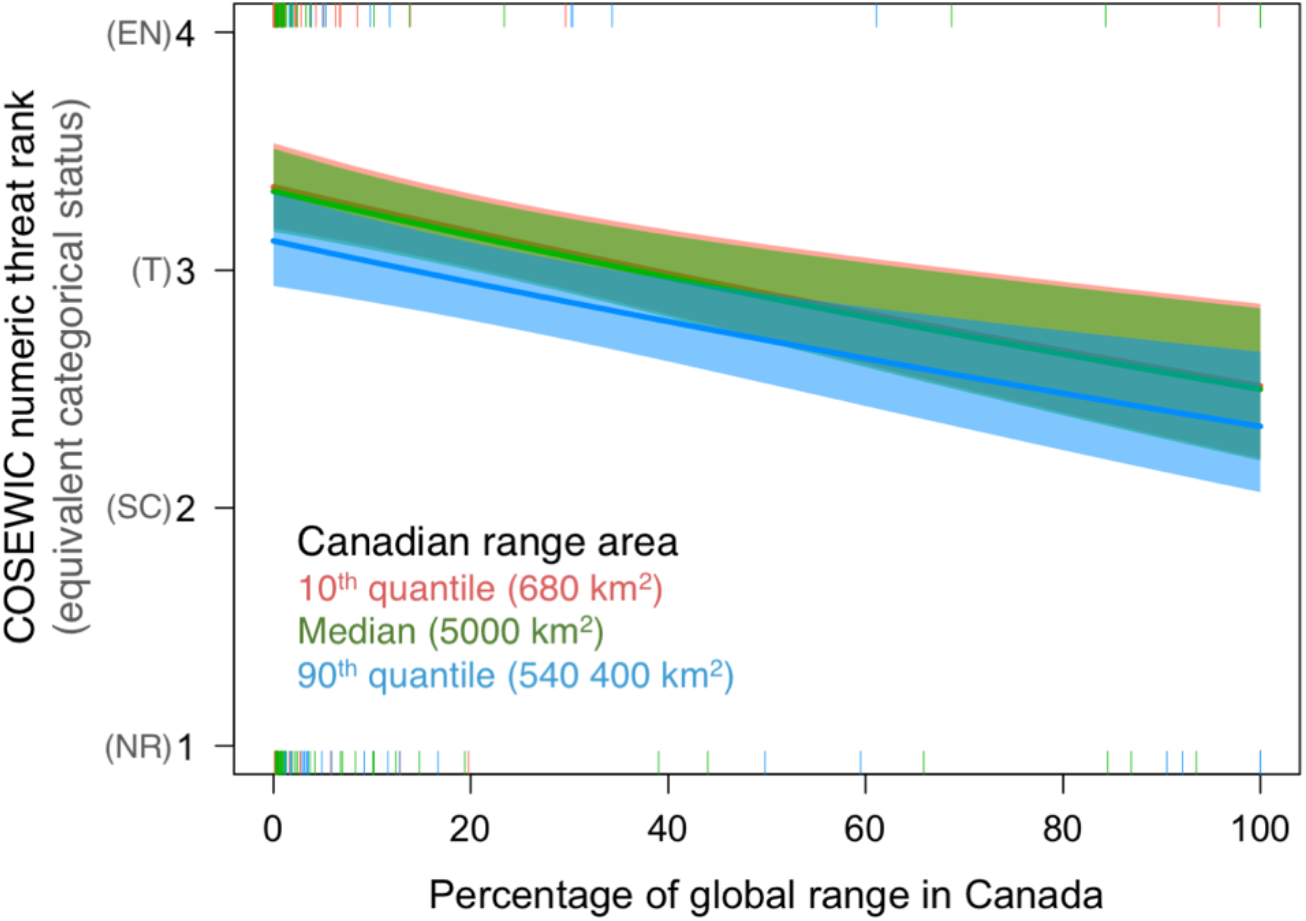
Being more peripheral (having a smaller % of global range) in Canada is associated with being more nationally threatened even after accounting for absolute range area in Canada (*Question 2*). QuassiPoisson GLM: numeric COSEWIC status ∼ Canadian range area + Canadian range fraction. Canadian range area is numeric, but is displayed categorically to help visualize patterns.

A colleague who served on COSEWIC’s vascular plants subcommittee pointed out that the way COSEWIC prioritizes and assesses taxa changed once the Species at Risk Act was passed in 2002, potentially reducing prioritization and ranking of peripheral populations. We tested this by re-running models in Question 2 with a categorical predictor denoting whether species had been assessed by COSEWIC before 2002. Taxa assessed before or after SARA did not vary in their global range area, Canadian range area, or fraction of their global range in Canada (Table S2). Taxa were designated as slightly more at risk after SARA passed (least squared mean numeric COSEWIC rank = 3.2, where 1=Not at risk and 4=Endangered), than before (mean = 2.5) potentially indicating more effective prioritization. The effect of range area or percent on COSEWIC status did not differ between taxa assessed before or after SARA (model with vs. without interactions: χ^2^_df=2_, *P* = 0.50).

**Table S2.** Reanalyzing models in Question 2 accounting for whether taxa were assessed before or after SARA passed in 2002.

| Response | Predictors in model | Predictor significance<br>$\chi^2_{df=1}, P$ |
| --- | --- | --- |
| Global range area | COSEWIC threat rank | 7.50, $P = 0.057$ |
| | Before/After SARA | 0.01, $P = 0.91$ |
| Canadian range area | COSEWIC threat rank | 21.1, $P < 0.001$ |
| | Before/After SARA | 3.3, $P = 0.070$ |
| % range in Canada | COSEWIC threat rank | 13.8, $P = 0.003$ |
| | Before/After SARA | 1.5, $P = 0.16$ |
| COSEWIC threat rank<br>(numeric) | Canadian range area | 11.7, $P < 0.0001$ |
| | Percent range in Canada | 4.7, $P < 0.031$ |
| | Before/After SARA | 17.2, $P < 0.0001$ |

### QUESTION 5

#### Changing definition of peripheral

Our main analyses for Q5 used 189 at-risk taxa for which we could quantify or estimate peripherality, defined as <20% of their global range in Canada (Table 1b). To test whether conclusions were sensitive to this definition of peripheral, we reran analyses using only at-risk taxa with a digitized range map and quantitative assessment of peripherality (*n* = 166), and again using a stricter threshold for ‘peripheral’ (≤10% of range in Canada). The effect of being peripheral generally remained non-significant, except for the number of studies that included Canadian populations (Table S3).

**Table S3.** Reanalysis of Question 5 with different definitions of peripheral. Note that Q5 considers only at-risk taxa. Cells give χ^2^_df=1_ test statistics for effect of peripheral (yes or no) and associated *P* values, df = 1 in all cases. Leftmost results column gives results reported in main manuscript, using all taxa for which peripherality could be quantified or estimated (Table 1b).

|  | Threshold for peripheral (% global range in Canada) |  |  |
| --- | --- | --- | --- |
|  | Taxa considered |  |  |
|  | N species (peripheral spp) |  |  |
| GLM<br>(error distribution) | <20%<br>COSEWIC map<br>n = 189 (159) | <20%<br>Digitized map<br>n = 166 (132) | $\leq 10\%$<br>Digitized map<br>n = 166 (120) |
| Likelihood studied anywhere<br>(qbinomial) | $\chi^2 = 1.73$<br>$P = 0.19$ | $\chi^2 = 1.71$<br>$P = 0.19$ | $\chi^2 = 0.02$<br>$P = 0.88$ |
| Studies per species<br>(negative binomial) | $\chi^2 = 0.21$<br>$P = 0.88$ | $\chi^2 = 0.26$<br>$P = 0.61$ | $\chi^2 = 1.10$<br>$P = 0.29$ |
| Likelihood studied in Canada<br>(qbinomial) | $\chi^2 = 0.02$<br>$P = 0.90$ | $\chi^2 = 0.12$<br>$P = 0.90$ | $\chi^2 = 1.32$<br>$P = 0.25$ |
| Studies per species in Canada<br>(negative binomial) | $\chi^2 = 2.72$<br>$P = 0.099$ | $\chi^2 = 3.66$<br>$P = 0.056$ | $\chi^2 = 6.97$<br>$P = 0.008$ |

## REFERENCES

Anderson, J., and Wadgymar, S. 2020. Climate change disrupts local adaptation and favours upslope migration. Ecol. Lett. 23(1): 181–192.

Bivand, R., and Rundel, C. 2018. rgeos: Interface to Geometry Engine - Open Source (’GEOS’). Available from https://cran.r-project.org/package=rgeos.

Boles, R.L., Lovett-Doust, J., and Lovett-Doust, L. 2000. Population genetic structure in green dragon (Arisaema dracontium, Araceae). Can. J. Bot. 77(10): 1401–1410. doi:10.1139/b99-089.

Breheny, P., and Burchett, W. 2017. Visualization of Regression Models Using visreg.

Brito, D., Ambal, R.G., Brooks, T., Silva, N.D., Foster, M., Hao, W., Hilton-Taylor, C., Paglia, A., Rodríguez, J.P., and Rodríguez, J.V. 2010. How similar are national red lists and the IUCN Red List? Biol. Conserv. 143(5): 1154–1158. doi:10.1016/j.biocon.2010.02.015.

Brown, J.H., Stevens, G.C., and Kaufman, D.M. 1996. The geographic range: size, shape, and internal structure. Annu. Rev. Ecol. Syst. 27: 597–623. doi:10.1146/annurev.ecolsys.27.1.597.

Budd, C., Zimmer, E., and Freeland, J.R. 2015. Conservation genetics of Magnolia acuminata, an endangered species in Canada: Can genetic diversity be maintained in fragmented, peripheral populations? Conserv. Genet. 16(6): 1359–1373. doi:10.1007/s10592-015-0746-9.

Bunnell, F.L., Campbell, R.W., and Squires, K.A. 2004. Conservation priorities for peripheral species: the example of British Columbia. Can. J. For. Res. 34(11): 2240–2247. doi:10.1139/x04-102.

Cameron, V., and Hargreaves, A.L. 2020. Spatial distribution and hotspots of mammals in Canada. bioRxiv: 2020.01.29.925461. doi:10.1101/2020.01.29.925461.

CCEA. 2016. Conservation Areas Reporting and Tracking System Procedures Manual and Database Schema Implementation version - 2016.: 43.

Chen, I.-C., Hill, J.K., Ohlemüller, R., Roy, D.B., and Thomas, C.D. 2011. Rapid range shifts of species associated with high levels of climate warming. Science 333(6045): 1024–6. doi:10.1126/science.1206432.

Ciotir, C., Yesson, C., and Freeland, J. 2013. The evolutionary history and conservation value of disjunct Bartonia paniculata subsp paniculata (Branched Bartonia) populations in Canada. Botany 91(9): 605–613. doi:10.1139/cjb-2013-0063.

Coristine, L.E., Jacob, A.L., Schuster, R., Ford, A., Woodley, S., Shiffman, D.S., Orihel, D., Baron, N.E., Bennett, N.J., Venter, O., Bittick, S.J., Favaro, B., Dey, C., Otto, S.P., Polfus, J.L., Nowlan, L., and Palen, W.J. 2018. Informing Canada’s commitment to biodiversity conservation: A science-based framework to help guide protected areas designation through Target 1 and beyond. Facets 3(1): 531–562. doi:10.1139/facets-2017-0102.

Coristine, L.E., and Kerr, J.T. 2011. Habitat loss, climate change, and emerging conservation challenges in Canada. Can. J. Zool. 89(5): 435–451. doi:10.1139/z11-023.

Eckert, C.G., Samis, K.E., and Lougheed, S.C. 2008. Genetic variation across species’ geographical ranges: the central–marginal hypothesis and beyond. Mol. Ecol. 17(5): 1170–1188. doi:10.1111/j.1365-294X.2007.03659.x.

Ellstrand, N.C., and Elam, D.R. 2003. Population Genetic Consequences of Small Population Size: Implications for Plant Conservation. Annu. Rev. Ecol. Syst. 24(1): 217–242. doi:10.1146/annurev.es.24.110193.001245.

Environment and Climate Change Canada. 2017. COSEWIC wildlife species assessment: candidate wildlife species. Available from https://www.canada.ca/en/environment-climate-change/services/committee-status-endangered-wildlife/wildlife-species-assessment-process-categories-guidelines/candidate.html [accessed 18 November 2019].

Freeman, B.G., Lee-Yaw, J.A.,, Sunday, J.M., and Hargreaves, A.L. 2018. Expanding, shifting and shrinking: The impact of global warming on species’ elevational distributions. Glob. Ecol. Biogeogr. 27(11): 1268–1276. doi:10.1111/geb.12774.

Gibson, S.Y., Van der Marel, R.C., and Starzomski, B.M. 2009. Climate change and conservation of leading-edge peripheral populations. Conserv. Biol. 23(6): 1369–1373. doi:10.1111/j.1523-1739.2009.01375.x.

Glass, W.R., Corkum, L.D., and Mandrak, N.E. 2017. Living on the edge: Traits of freshwater fish species at risk in Canada. Aquat. Conserv. Mar. Freshw. Ecosyst. 27(5): 938–945. doi:10.1002/aqc.2781.

Godt, M.J.W., Caplow, F., and Hamrick, J.L. 2005. Allozyme diversity in the federally threatened golden paintbrush, Castilleja levisecta (Scrophulariaceae). Conserv. Genet. 6(1): 87–99. doi:10.1007/s10592-004-7746-5.

Government of Canada. 2019. Canada Nature Fund. Available from https://www.canada.ca/en/environment-climate-change/services/nature-legacy/fund.html [accessed 16 June 2019].

Halbritter, A.H., Alexander, J.M., Edwards, P.J., and Billeter, R. 2013. How comparable are species distributions along elevational and latitudinal climate gradients? Glob. Ecol. Biogeogr. 22(11): 1228–1237. doi:10.1111/geb.12066.

Hampe, A., and Petit, R.J. 2005. Conserving biodiversity under climate change: The rear edge matters. Ecol. Lett. 8(5): 461–467. doi:10.1111/j.1461-0248.2005.00739.x.

Hargreaves, A.L., Bailey, S.F., and Laird, R.A. 2015. Fitness declines towards range limits and local adaptation to climate affect dispersal evolution during climate-induced range shifts. J. Evol. Biol. 28(8): 1489–1501. doi:10.1111/jeb.12669.

Hargreaves, A.L., and Eckert, C.G. 2019. Local adaptation primes cold-edge populations for range expansion but not warming-induced range shifts. Ecol. Lett. 22(1): 78–88. doi:10.1111/ele.13169.

Hargreaves, A.L., Samis, K.E., and Eckert, C.G. 2014. Are Species’ Range Limits Simply Niche Limits Writ Large? A Review of Transplant Experiments beyond the Range. Am. Nat. 183(2): 157–173. The University of Chicago PressThe American Society of Naturalists. doi:10.1086/674525.

Hengeveld, R., and Haeck, J. 1982. The distribution of abundance. J. Biogeogr. 9(4): 303–316.

Hoban, S.M., Borkowski, D.S., Brosi, S.L., McCleary, T.S., Thompson, L.M., McLachlan, J.S., Pereira, M.A., Schlarbaum, S.E., and Romero-Severson, J. 2010. Range-wide distribution of genetic diversity in the North American tree Juglans cinerea: a product of range shifts, not ecological marginality or recent population decline. Mol. Ecol. 19(22): 4876–4891. doi:10.1111/j.1365-294X.2010.04834.x.

Hunter, M.I., and Hutchinson, A. 1994. The Virtues and Shortcomings of Parochialism: Conserving Species That Are Locally Rare, but Globally Common. Conserv. Biol. 8(4): 1163–1165.

Komonen, A. 2007. Are we conserving peripheral populations? An analysis of range structure of longhorn beetles (Coleoptera: Cerambycidae) in Finland. J. Insect Conserv. 11(3): 281–285. doi:10.1007/s10841-006-9043-8.

Lee-Yaw, J.A.,, Kharouba, H.M., Bontrager, M., Mahony, C., Csergő, A.M., Noreen, A.M.E., Li, Q., Schuster, R., and Angert, A.L. 2016. A synthesis of transplant experiments and ecological niche models suggests that range limits are often niche limits. Ecol. Lett. 19(6): 710–722. doi:10.1111/ele.12604.

Lenth, R.V. 2016. Least-Squares Means: The R Package lsmeans. doi:10.18637/jss.v069.i01.

Lesica, P., Adams, B., and Smith, C.T. 2016. Can physiographic regions substitute for genetically-determined conservation units? A case study with the threatened plant, Silene spaldingii. Conserv. Genet. 17(5): 1041–1054. doi:10.1007/s10592-016-0842-5.

Lesica, P., and Allendorf, F.W. 1995. When Are Peripheral Populations Valuable for Conservation? Conserv. Biol. 9(4): 753–760.

McKie, D. 2016. David McKie | Your home for data journalism. Available from http://www.davidmckie.com/ [accessed 22 December 2018].

MDEL. 2016. Registre des aires protégées au Québec.

de Medeiros, C.M., Hernández-Lambraño, R.E., Ribeiro, K.A.F., and Sánchez Agudo, J.Á. 2018. Living on the edge: do central and marginal populations of plants differ in habitat suitability? Plant Ecol. 219(9): 1029–1043. doi:10.1007/s11258-018-0855-x.

Mehrhoff, L.A. 1989. Reproductive vigor and environmental factors in populations of an endangered North American orchid, Isotria medeoloides (Pursh) rafinesque. Biol. Conserv. 47(4): 281–296. doi:10.1016/0006-3207(89)90071-2.

Mymudes, M.S., and Les, D.H. 1993. Morphological and Genetic Variability in Plantago cordata (Plantaginaceae), a Threatened Aquatic Plant. Am. J. Bot. 80(3): 351–359. doi:10.2307/2445359.

NatureServe. 2018. NatureServe Explorer: NatureServe Conservation Status. Available from http://explorer.natureserve.org/ranking.htm [accessed 17 November 2019].

Phillips, B.L., Brown, G.P., and Shine, R. 2010. Evolutionarily accelerated invasions: The rate of dispersal evolves upwards during the range advance of cane toads. J. Evol. Biol. 23(12): 2595–2601. doi:10.1111/j.1420-9101.2010.02118.x.

Pironon, S., Papuga, G., Villellas, J., Angert, A.L., García, M.B., and Thompson, J.D. 2017. Geographic variation in genetic and demographic performance: new insights from an old biogeographical paradigm. Biol. Rev. 92(4): 1877–1909. doi:10.1111/brv.12313.

Pironon, S., Villellas, J., Morris, W.F., Doak, D.F., and García, M.B. 2015. Do geographic, climatic or historical ranges differentiate the performance of central versus peripheral populations? Glob. Ecol. Biogeogr. 24(6): 611–620. doi:10.1111/geb.12263.

QGIS Development Team. 2018. QGIS Geographic Information System. Available from http://qgis.osgeo.org.

R Core Team. 2017. R: A language and environment for statistical computing. Vienna. Available from https://www.r-project.org/.

Raven, P. 1987. Scope of the plant conservation problem world-wide in Bramwell D, editor. In Proceedings of an International Conference on botanic gardens and the world conservation strategy. London: Academic Press., Las Palmas de Gran Canaria.

Sagarin, R.D., Gaines, S.D., and Gaylord, B. 2006. Moving beyond assumptions to understand abundance distributions across the ranges of species. Trends Ecol. Evol. 21(9): 524–530. doi:10.1016/j.tree.2006.06.008.

SARA. 2002. Species at Risk Act.

Schemske, D.W., Husband, B.C., Ruckelshaus, M.H., Goodwillie, C., Parker, I.M., and Bishop, J.G. 1994. Evaluating approaches to the conservation of rare and endangered plants. Ecology 75(3): 584–606.

Sexton, J.P., McIntyre, P.J., Angert, A.L., and Rice, K.J. 2009. Evolution and Ecology of Species Range Limits. Annu. Rev. Ecol. Evol. Syst. 40(1): 415–436. doi:10.1146/annurev.ecolsys.110308.120317.

Sexton, J.P., Strauss, S.Y., and Rice, K.J. 2011. Gene flow increases fitness at the warm edge of a species’ range. Proc. Natl. Acad. Sci. 108(28): 11704–11709. doi:10.1073/pnas.1100404108.

Shaw, R.G., and Etterson, J.R. 2012. Rapid climate change and the rate of adaptation: insight from experimental quantitative genetics. New Phytol. 195(4): 752–765. doi:10.1111/j.1469-8137.2012.04230.x.

Statistics Canada. 2011. Census Dictionary.

Statistics Canada. 2016. Geographic Attribute File, Reference Guide.

Steen, D.A., and Barrett, K. 2015. Should states in the USA value species at the edge of their geographic range? J. Wildl. Manag. 79(6): 872–876. doi:10.1002/jwmg.897.

Thakur, M., Schättin, E.W., and McShea, W.J. 2018. Globally common, locally rare: revisiting disregarded genetic diversity for conservation planning of widespread species. Biodivers. Conserv. 27(11): 1–5. doi:10.1007/s10531-018-1579-x.

Thomas, C.D., Cameron, A., Green, R.E., Bakkenes, M., Beaumont, L.J., Collingham, Y.C., Erasmus, B.F.N., de Siqueira, M.F., Grainger, A., Hannah, L., Hughes, L., Huntley, B., van Jaarsveld, A.S., Midgley, G.F., Miles, L., Ortega-Huerta, M.A.,, Townsend Peterson, A., Phillips, O.L., and Williams, S.E. 2004. Extinction risk from climate change. Nature 427(6970): 145–148. doi:10.1038/nature02121.

Usery, E.L., and Seong, J.-C. 2000. A Comparison of Equal-Area Map Projections for Regional and Global Raster Data. PhD Thesis. Available from http://carto-research.er.usgs.gov/projection/pdf/nmdrs.usery.prn.pdf.

Villellas, J., Morris, W.F., and Garcia, M.B. 2013. Variation in stochastic demography between and within central and peripheral regions in a widespread short-lived herb. Ecology 94(6): 1378–1388. doi:10.1890/12-1163.1.

Yakimowski, S.B., and Eckert, C.G. 2007. Threatened peripheral populations in context: Geographical variation in population frequency and size and sexual reproduction in a clonal woody shrub. Conserv. Biol. 21(3): 811–822. doi:10.1111/j.1523-1739.2007.00684.x.

Yakimowski, S.B., and Eckert, C.G. 2008. Populations do not become less genetically diverse or more differentiated towards the northern limit of the geographical range in clonal Vaccinium stamineum (Ericaceae). New Phytol. 180(2): 534–544. doi:10.1111/j.1469-8137.2008.02582.x.

Yang, J., Lovett-Doust, J., and Lovett-Doust, L. 1999. Seed germination patterns in green dragon (Arisaema dracontium, Araceae). Am. J. Bot. 86(8): 1160–1167. doi:10.2307/2656980.

Yeaman, S., Hodgins, K.A., Lotterhos, K.E., Suren, H., Nadeau, S., Degner, J.C., Nurkowski, K.A., Smets, P., Wang, T., Gray, L.K., Liepe, K.J., Hamann, A., Holliday, J.A., Whitlock, M.C., Rieseberg, L.H., and Aitken, S.N. 2016. Convergent local adaptation to climate in distantly related conifers. Science 353(6306): 1431–1433. doi:10.1126/science.aaf7812.

